# Stochastic Modeling of Tinnitus Loudness

**DOI:** 10.1101/2023.02.09.527783

**Authors:** Sangyeop Kwak, Daehee Lee, Sungshin Jang, Songhwa Kim, Sunghwan Kim, Woojin Doo, Eunyee Kwak

**Author notes:** **Correspondence:** Sangyeop Kwak.

## Abstract

There has been no study on the relationship between chronic tinnitus and harmonic templates. Harmonic templates are harmonically structured receptive fields in the auditory system in which all frequency components are integer multiples of a common fundamental frequency (F_0_). In this study, data from 19 harmonic templates from each of 196 chronic tinnitus patients were analyzed and mathematical modeling was performed to quantify the loudness of chronic tinnitus. High-resolution hearing threshold data were obtained by algorithmic pure tone audiometry (PTA) conducting automated PTA at 134 frequency bands with 1/24 octave resolution from 250 Hz to 12,000 Hz. The result showed that there is an intriguing relationship between the auditory instability of harmonic templates and simplified tinnitus severity score (STSS). This study provides several mathematical models to estimate tinnitus severity and the precise quantification of the loudness of chronic tinnitus. Our computational models and analysis of the behavioral hearing threshold fine structure suggest that the cause of severe chronic tinnitus could be a severe disparity between different temporal capacities of each neural oscillator in a certain harmonic template.

## Introduction

Chronic tinnitus as a perceived sound has acoustic characteristics. The timbre of tinnitus is characterized by either the simplicity or complexity of the acoustic spectrum in which a salient tinnitus frequency region is defined as a function of frequency, intensity and time. Timbre and loudness are key auditory percepts of sound. Although most studies on the temporal fine structure of the auditory system have focused on the pitch of the sound, the success or failure rate of auditory phase locking in the microsecond time domain probably underlies different aspects in the neural gain control mechanism for the perception of timbre and loudness^1, 2, 3^.

The existing practice for matching the pitch and loudness of a certain tinnitus relies on the listening tasks with a single pure tone or a narrow-band noise. Although there are a number of established methods for tinnitus pitch and loudness matching and tinnitus surveys^4, 5^, the reliability of those methods for quantifying the severity of chronic tinnitus is still questionable^6^. There could be many types of tinnitus spectra according to the deprived frequency regions in the hearing threshold fine structure. In tinnitus taxonomy, tinnitus is classified into five types according to its acoustic spectrum as simple tone, simple noise, complex tone, complex noise, and a combination of tone and noise^7^. Thus, there could be 10 types of chronic tinnitus according to whether tinnitus occurs unilaterally or bilaterally, as shown in Figure 1^7^.

**Figure 1.**
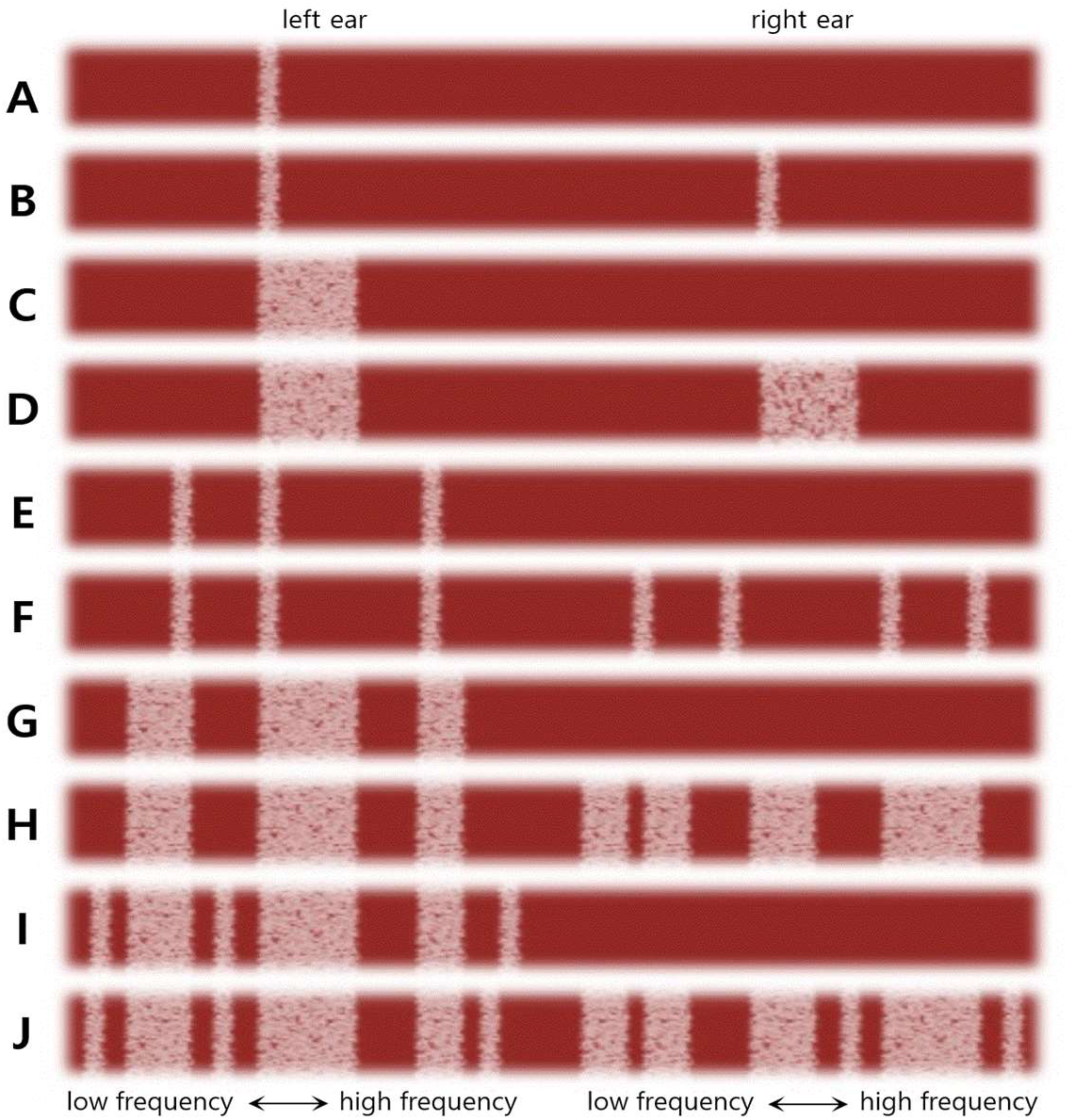
Simulated acoustic spectra of 10 types of chronic tinnitus. (A) Unilateral simple tone, (B) bilateral simple tone, (C) unilateral simple noise, (D) bilateral simple noise, (E) unilateral complex tone, (F) bilateral complex tone, (G) unilateral complex noise, (H) bilateral complex noise, (I) unilateral combination of tone and noise, and (J) bilateral combination of tone and noise.

A recent study on missed hearing loss in tinnitus patients with normal audiograms found that there were many cases of sharply notched hearing loss in normal audiograms and that most notches were at tinnitus frequencies^8^. However, most studies and clinical practice related to the pitch and loudness of chronic tinnitus have focused on only four to 10 frequencies from traditional pure tone audiometry (PTA) using a few pure tones or a couple of narrow-band noises for their pitch and loudness matching tasks. The methodology in the present study focused on the instability of harmonic templates provides a comprehensive insight into the correlation between the behavioral hearing threshold fine structure and the severity of chronic tinnitus. Harmonic templates are harmonically structured receptive fields in the auditory system in which all frequency components are integer multiples of a common fundamental frequency (F_0_)^9, 10^.

There has been no study reporting systematic measurements of the tinnitus spectrum showing its pitch and loudness information simultaneously. The frequencies employed in this study are beyond the conventional frequency profile in PTA which has been widely accepted in current clinical practice. We took a novel approach with 43 equal temperament frequencies in the range from 1,047 Hz to 11,840 Hz to grasp the chronic tinnitus spectrum at the level of the auditory threshold fine structure containing its footprint of salient tinnitus information. Spectrally salient regions of chronic tinnitus do not necessarily follow the four to 10 frequencies adopted by existing audiometric standards^8, 11^.

This study investigated a way to quantify the severity of chronic tinnitus using two concepts. One was the auditory instability in a certain harmonic template. The other was a three-dimensional (3D) Cartesian coordinate in which the X-axis represented the fundamental frequency of harmonic templates, the Y-axis showed the auditory capacity of the harmonic templates, and the Z-axis indicated the level of auditory instability at a certain harmonic template. The auditory capacity as a subsidiary concept to formulate the concept of auditory instability was adopted to scrutinize the level of auditory instability in each of 19 harmonic templates. To confirm the feasibility of the first concept, we collected and analyzed real human data on the behavioral hearing threshold fine structure acquired from chronic tinnitus patients.

We hypothesize that auditory instability at a certain harmonic template is a key parameter of the appropriate model for the loudness estimation of chronic tinnitus. Along with our endeavor to quantify the loudness level of chronic tinnitus, this study proposes a model based on two notions. The first notion was that the unstable harmonic template might be the cause of the lower success rate of auditory phase locking resulting in the higher loudness level of chronic tinnitus. The second notion was that the stability of the harmonic template contributes to the reduced severity of chronic tinnitus. It is noteworthy that the most stable condition in hearing could indicate not only perfectly normal hearing but also complete hearing loss.

## Materials and Methods

### Stochastic representation of the auditory capacity and instability of harmonic templates

This study speculates on the neural network of the active auditory system as a stochastic vector field. We defined the harmonic template in the human active auditory system as ‘Fz’ or ‘fz’. The auditory capacity and instability of the harmonic templates were evaluated by the function(*f*) of Fz and fz. Both ‘F_0_’ and ‘f_0_’ represented a fundamental frequency of a certain Fz. Complex impedance was indicated by the lowercase letter ‘z’. The auditory capacity of a certain Fz was indicated by μ(fz) which was the mean (μ) threshold in the dB hearing level (dBHL) of f_n_ (n = 0, 1, 2, 3) truncated at the fourth harmonics of a certain f_0_. The term μ(fz) was defined in this study as the mean value of mu (μ) construed by at least 24 behavioral hearing thresholds gained at f_0_, f_1_, f_2_, and f_3_ in both ears (4 frequencies x 2 ears x 3 tests).

The standard deviation (σ) value was employed to evaluate the auditory instability at a certain μ(fz). The term σ(f_n_) was defined as a function of the sigma (σ) level of the standard deviation induced by at least 24 behavioral hearing thresholds gained at f_0_, f_1_, f_2_, and f_3_ in both ears (4 frequencies x 2 ears x 3 tests). To confirm the estimated data from the model, we analyzed real hearing data for the 19 harmonic templates from 01(Fz) to 19(Fz), which included four frequency entries in each row vector of f_0_(Fz) as shown in Table 1.

**Table 1.** Row vector of f0(Fz)

| Fz | F<sub>0</sub> | Row vector of $f_0(Fz)$, $f_0$ = fundamental frequency in Hz<br>$f_1 = 2f_0, f_2 = 3f_0, f_3 = 4f_0$ |
| --- | --- | --- |
| 19 | 2960 | [ 2960 5920 8870 11840 ] |
| 18 | 2794 | [ 2794 5588 8372 11175 ] |
| 17 | 2637 | [ 2637 5274 7902 10548 ] |
| 16 | 2489 | [ 2489 4978 7459 9956 ] |
| 15 | 2349 | [ 2349 4699 7040 9397 ] |
| 14 | 2217 | [ 2217 4435 6645 8870 ] |
| 13 | 2093 | [ 2093 4186 6272 8372 ] |
| 12 | 1976 | [ 1976 3951 5920 7902 ] |
| 11 | 1865 | [ 1865 3729 5588 7459 ] |
| 10 | 1760 | [ 1760 3520 5274 7040 ] |
| 9 | 1661 | [ 1661 3322 4978 6645 ] |
| 8 | 1568 | [ 1568 3136 4699 6272 ] |
| 7 | 1480 | [ 1480 2960 4435 5920 ] |
| 6 | 1397 | [ 1397 2794 4186 5588 ] |
| 5 | 1319 | [ 1319 2637 3951 5274 ] |
| 4 | 1245 | [ 1245 2489 3729 4978 ] |
| 3 | 1175 | [ 1175 2349 3520 4699 ] |
| 2 | 1109 | [ 1109 2217 3322 4435 ] |
| 1 | 1047 | [ 1047 2093 3136 4186 ] |

### The Gaussian integral to evaluate tinnitus severity

#### The loudness estimation of simple tinnitus

We defined simple tinnitus as either single sinusoidal tone or single narrow-band noise. All kinds of simple tinnitus should have only their own spectral peaks in a single auditory filter. The passive trajectory *L_p_* of the loudness of simple tinnitus in the passive auditory system was assumed to follow a sinusoidal arc against the behavioral hearing threshold *y* from the 20 to the 90 dBHL in the passive auditory system. The loudness of the auditory input below the 20 dBHL or above the 90 dBHL may be affected by efferent induced damping and recruitment^12, 13, 14, 15^. Thus, additional correction functions might be needed to quantify simple tinnitus occurring within the behavioral hearing thresholds below the 20 dBHL or above the 90 dBHL. The passive trajectory *L_p_* of the loudness of simple tinnitus in the passive auditory system was defined as follows:

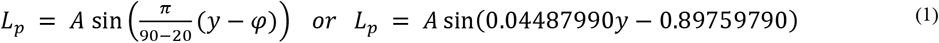

where *A* is the amplitude and *φ* is the phase. Unlike tinnitus loudness in the passive auditory system, it was assumed that the active trajectory *L_a_* of the loudness of simple tinnitus in the human active auditory system followed a Gaussian curve against the behavioral hearing threshold *y* from the −10 to the 120 dBHL. We referred to this as Gaussian correction in simple tinnitus reflecting efferent-induced damping and recruitment in the human active auditory system. The formula to define the active trajectory *L_a_* of simple tinnitus loudness in human active harmonic templates was:

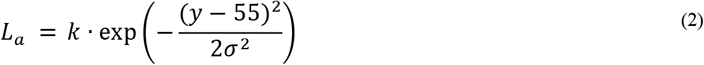

where *k* is the coefficient to determine a peak or a plateau in the Gaussian trajectory, and σ is the standard deviation of the behavioral hearing thresholds in a certain auditory unit.

#### The z-weighted loudness estimation of chronic tinnitus

Tinnitus in the human active auditory system is a stochastic neural aspect in the nonlinear auditory neural circuit. Various types of tinnitus spectra are accounted for as auditory percepts using linear and nonlinear encoding in deep neural networks^16^. We defined the complex impedance in the active auditory system by the lowercase letter z. The temporal fine structure in the active auditory circuit might be constrained by neural AC/DC impedance control. Z-weighted correction in the present study was used to closely examine the stochastic distance between both levels of auditory instability σ(f_n_) and the loudness of chronic tinnitus determined by neural AC/DC impedance associated with tinnitus occurrence. The level of σ(f_n_) against the auditory capacity μ(fz) in dBHL was represented by the height in the 2D Gaussian trajectory with a unit of dBzHL, which was proposed in this study for the first time. An unknown neural AC/DC impedance associated with tinnitus occurrence in the human active auditory system might determine the distance between the actually perceived loudness of tinnitus and the level of σ(f_n_) at a certain Fz. In this context, a σ(f_n_) plays an important role as a stochastic parameter to evaluate the perceived loudness level of chronic tinnitus. According to our computational model, the actually perceived loudness level of simple tinnitus in the active auditory system was predicted to be equivalent to the z-weighted Gaussian integral. The z-weighting method is not different from the weighted evaluation of a Gaussian integral to estimate the loudness level of either simple, complex, or combined tinnitus in human active harmonic templates. The timbre of tinnitus in most chronic tinnitus patients is considered to have complex or combined spectra in which the loudness levels of a number of simple tinnitus components are summed across individual auditory filters. If the tinnitus loudness is summed across a number of individual auditory filters, then it is assumed that the active trajectory *L_c_* of the loudness of chronic tinnitus follows the horizontally flipped Rayleigh curve derived from the z-weighted Gaussian integral in human active harmonic templates against the behavioral hearing threshold *y* defined as follows:

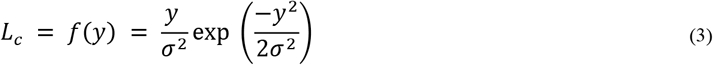

Formula (3) is a function of the normal Rayleigh distribution before horizontal flipping^17, 18^.

We considered complex tinnitus in human active harmonic templates as a stochastic aspect determined by the complex impedance z by which the loudness of complex tinnitus was constituted at a level beyond the simple sum of the loudness levels of individual simple tinnitus components. There are numerous types of complex tinnitus including the combined type where both types of tonal and noise tinnitus coexist across a number of individual auditory filters. If a tinnitus spectrum is assumed to have multiple salient regions at different auditory filters, the 3D Gaussian curvature might be more useful for estimating the loudness of chronic tinnitus not only against the Y-axis representing the auditory capacity but also against the X-axis representing the fundamental frequency of harmonic templates. The level of σ(f_n_) in complex tinnitus was represented by the height in the 3D trajectory of Gaussian curvature in dBzHL. The actually perceived loudness level of complex tinnitus was assumed to follow the 3D trajectory of Gaussian curvature against the Fz-axis in Hz and the μ(fz)-axis in dBHL. The perceived loudness level of complex tinnitus can also be defined as follows:

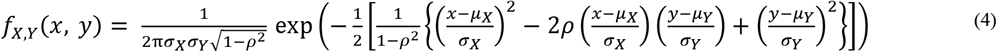

where *μ_x_* indicates the mean of one Gaussian curve against the X-axis in Hz from −19(Fz) to 19(Fz) as a function of the fundamental frequency of Fz, *μ_Y_* indicates the mean of the other Gaussian curve against the Y-axis of μ(fz) from the −10 to the 120 dBHL, and *ρ* is 0 because both X and Y variables are independent.

### Algorithmic pure tone audiometry (PTA) and hearing thresholds with 1/12 octave frequency resolution

We performed a retrospective study on anonymized fine-grained audiometric data from tinnitus patients who came to the clinics for medical examination. We did not recruit ‘human participants’ or ‘human subjects’ for our retrospective study. The ‘patients’ reported in the study were general tinnitus patients who visited clinics. Therefore, no ethical approval was needed in Korea for our retrospective data mining and computational work. A total of 50,568 behavioral hearing thresholds data were collected from 17 medical clinics where the algorithmic PTA was adopted for high-resolution audiometry and tinnitus diagnosis. Figure 2 shows the average audiogram of 196 chronic tinnitus patients tested by the algorithmic PTA with 1/12 octave resolution from 250 Hz to 12,000 Hz. The algorithmic PTA is a Korea Food and Drug Administration (KFDA)-approved class 2 software as a medical device (SaMD; approval number 05-945) which is the first computerized audiometer conducting automated PTA at 134 frequency bands with 1/24 octave resolution from 250 Hz to 12,000 Hz^19^. The raw data of 196 chronic tinnitus patients were analyzed in the present study. Each datum obtained from each patient consisted of 43 behavioral hearing thresholds in dBHL at 43 frequencies in Hz with 1/12 octave resolution from 1,047 Hz to 11,840 Hz in each ear, i.e., 86 hearing thresholds in both ears.

**Figure 2.**
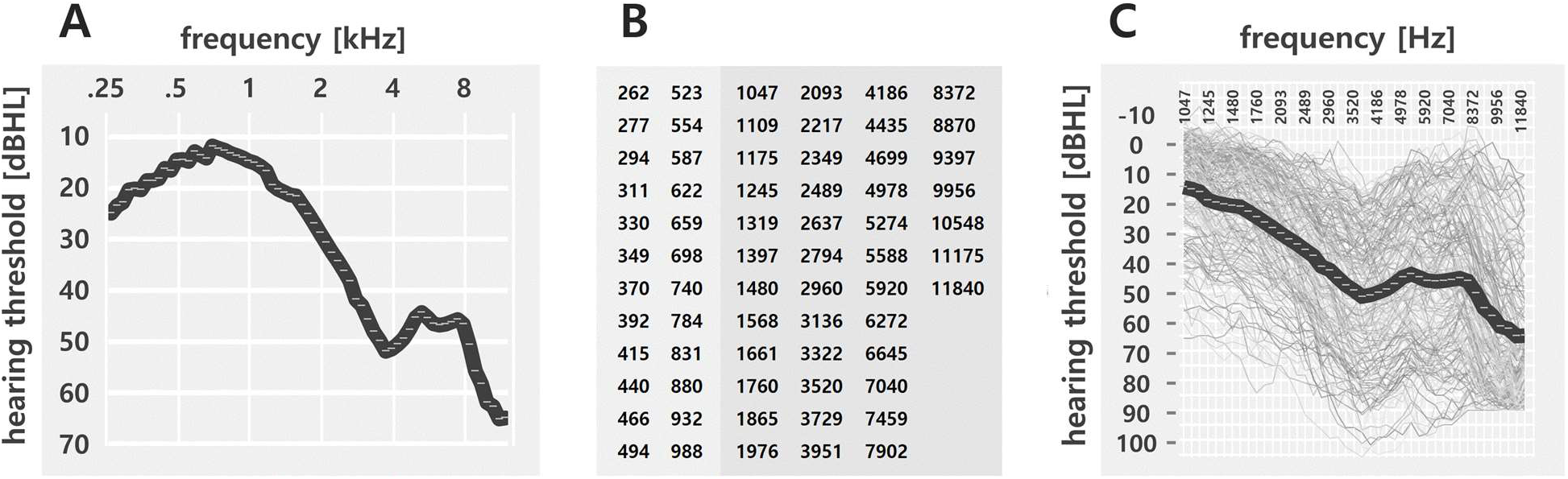
(A) Average audiogram of 196 chronic tinnitus patients with 1/12 octave resolution from 250 Hz to 12,000 Hz. (B) A profile of 67 frequencies tested by algorithmic PTA. The frequency profile is derived from the formula 2.92 (*f* x 1.05946 – *f*) based on ISO 16:1975 Acoustics - Standard tuning frequency (standard musical pitch). The average turnaround time for 196 chronic tinnitus patients to complete 1/12 octave resolution audiometry was 935 seconds (15’ 35”) from 250 Hz to 12,000 Hz in both ears. (C) The 196 audiograms at 43 frequencies with 1/12 octave resolution between 1 kHz and 12 kHz. The bold line is the average audiogram of 196 chronic tinnitus patients. The average turnaround time was 600 seconds (10’) for both ears to complete 1/12 octave resolution audiometry at 43 frequencies from 1,047 Hz to 11,840 Hz.

### Three-dimensional representation to integrate the auditory capacity and instability of the harmonic templates

We employed a 3D Cartesian coordinate system to represent the auditory capacity and instability at a certain Fz. The fundamental frequency of Fz was indicated on the X-axis as a function of frequency in Hz. The auditory capacity at a certain Fz was indicated on the Y-axis as the dBHL. A total of 3,724 mu (μ) values were acquired from 19 harmonic templates from each of 196 chronic tinnitus patients, as shown in the section A of Table 2. The level of auditory instability at a certain Fz was indicated on the Z-axis as a function of the dB hearing ‘z’ level (dBzHL). The dBzHL unit was employed in this study for the first time to indicate the level of auditory instability at a certain Fz. This novel unit differed from the dB sensation level (dBSL) widely used in the tinnitus loudness matching test. The origin of the dBzHL was based on the unknown stochastic impedance in the active harmonic templates correlated with the standard deviation (σ) induced by at least 24 behavioral hearing threshold entries in an Fz matrix (4 frequencies x 2 ears x 3 tests). In the present study, a total of 3,724 sigma (σ) levels from 01(Fz) to 19(Fz) were obtained from each of 196 chronic tinnitus patients, as shown in the section B of Table 2. Figure 3 shows 196 colored result lines representing auditory capacity μ(fz) against the 19 fundamental frequencies from 01(Fz) to 19(Fz). The average of the 196 result lines is indicated by 19 black circles. Its 2D representation was expressed in 3D Cartesian coordinates in this study to analyze and discuss the correlation between the level of σ(f_n_) and tinnitus severity in chronic tinnitus patients, as shown in Figure 3B and Figure 3C.

**Table 2.**
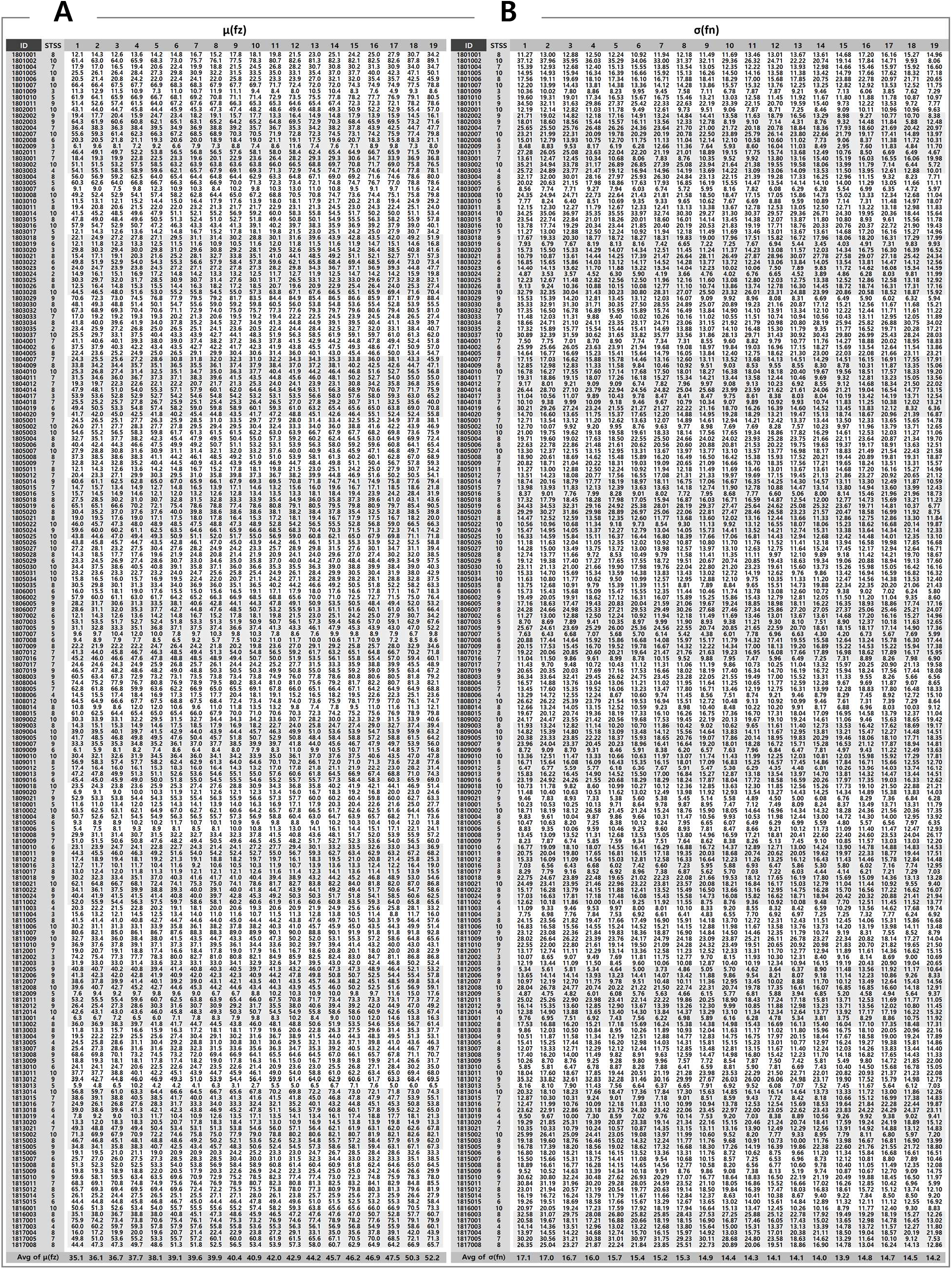
(A) Results of the level of STSS and 3,724 μ(fz) values of 196 chronic tinnitus patients. (B) Results of the level of STSS and 3,724 σ(fn) values of 196 chronic tinnitus patients.

**Figure 3.**
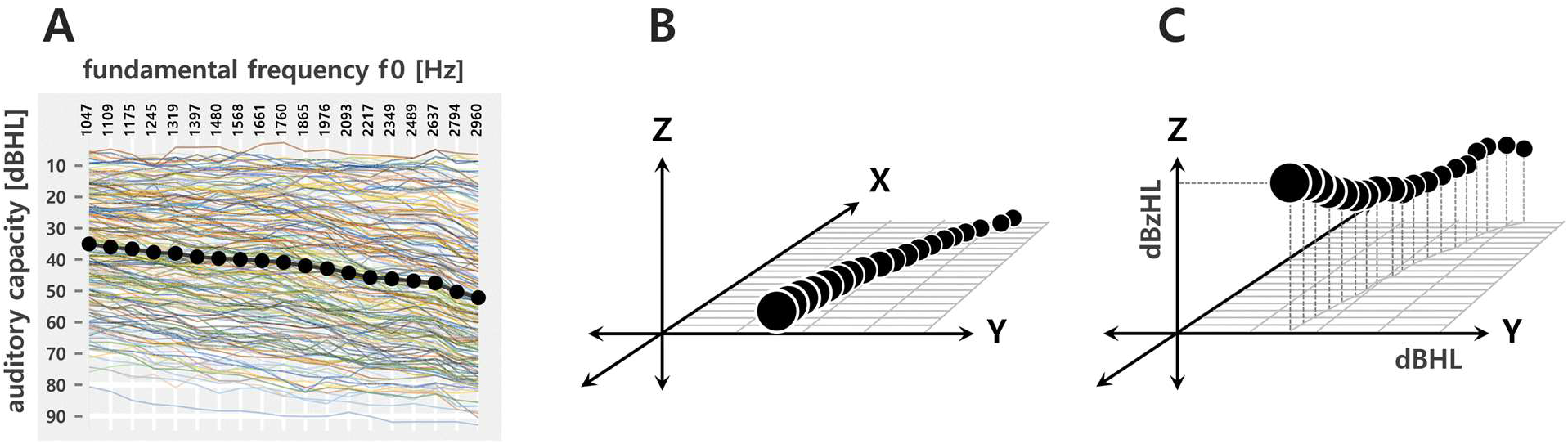
(A) A total of 196 result lines for the auditory capacity μ(fz) in different colors against 19 fundamental frequencies from 01(Fz) to 19(Fz). Each of 3,724 values of μ(fz) was considered the mean (μ) of the thresholds in dBHL of fn (n = 0, 1, 2, 3) truncated at the fourth harmonics of a certain f0. A set of 19 harmonic templates from 01(Fz) to 19(Fz) was equivalent to a similar set from 1,047(Fz) to 2,960(Fz). The 19 black circles denote an average of 196 result lines. The raw data are shown in the section A of Table 2. (B) An example of the 3D representation of μ(fz) on the Y-axis against the 19 fundamental frequencies of Fz on the X-axis. (C) An example of the 3D representation of σ(fn) on the Z-axis against the μ(fz) Y-axis and against the 19 fundamental frequencies of Fz on the X-axis. The raw data are shown in the section B of Table 2.

### Simplified tinnitus severity score (STSS)

A total of 196 chronic tinnitus patients were asked to choose one of the following five options of severe (5), moderate (4), mild (3), barely audible (2), and inaudible (1) to estimate tinnitus loudness. The question was “how is your tinnitus volume now?” To obtain a simplified tinnitus severity score (STSS), the clinicians asked this question twice at an interval of 4 weeks. A summed score of 10 consisted of severe (5) + severe (5). A summed score of 9 consisted of either severe (5) + moderate (4) or its inverse. A summed score of 8 might have one of the following three combinations: severe (5) + mild (3), mild (3) + severe (5), and moderate (4) + moderate (4). A summed score of 7 might have one of the following four combinations: severe (5) + barely audible (2), barely audible (2) + severe (5), moderate (4) + mild (3), and mild (3) + moderate (4). A summed score of 6 might have one of the following five combinations: severe (5) + inaudible (1), inaudible (1) + severe (5), moderate (4) + barely audible (2), barely audible (2) + moderate (4), and mild (3) + mild (3). A summed sore of 5 might have one of the four combinations: moderate (4) + inaudible (1), inaudible (1) + moderate (4), mild (3) + barely audible (2), and barely audible (2) + mild (3). A summed score of 4 might have one of the following three combinations: mild (3) + inaudible (1), inaudible (1) + mild (3), and barely audible (2) + barely audible (2). A summed score of 3 might have one of the following two combinations: barely audible (2) + inaudible (1) and inaudible (1) + barely audible (2). A summed score of 2 consisted of inaudible (1) + inaudible (1).

## Results

### Representation of auditory instability σ(fn) and simplified tinnitus severity score (STSS)

Different patterns of colors in the 19 representations in Figure 4A show different levels of σ(fn) against auditory capacity μ(fz) in each of the 19 harmonic templates (Fz). The four colors represent different ranges of σ(fn) levels along with 196 individual values in μ(fz) at a certain Fz. The nonlinear intervals between the small dots beside the colored bars indicate a 10-dB difference in μ(fz). The most unstable area tended to gather around the 50~60 dBHL in all of the 19 harmonic templates. The summed representation of the 19 bars and the normalized results are shown in Figure 4B. The different intervals between the black dots in the summed representation are due to different sample sizes of the μ(fz) values. A normalized representation with 200 equivalent intervals is shown in 0.5-dB steps from the 0 dBHL to the 100 dBHL. The summed result of the STSS levels (N = 3,724) against μ(fz) is shown in Figure 4C. The STSS levels of 196 patients were allocated equally into each of 19 harmonic templates of the 196 chronic tinnitus patients. A normalized representation of STSS is also shown with 200 equivalent intervals in 0.5-dB steps from the 0 dBHL to the 100 dBHL.

**Figure 4.**
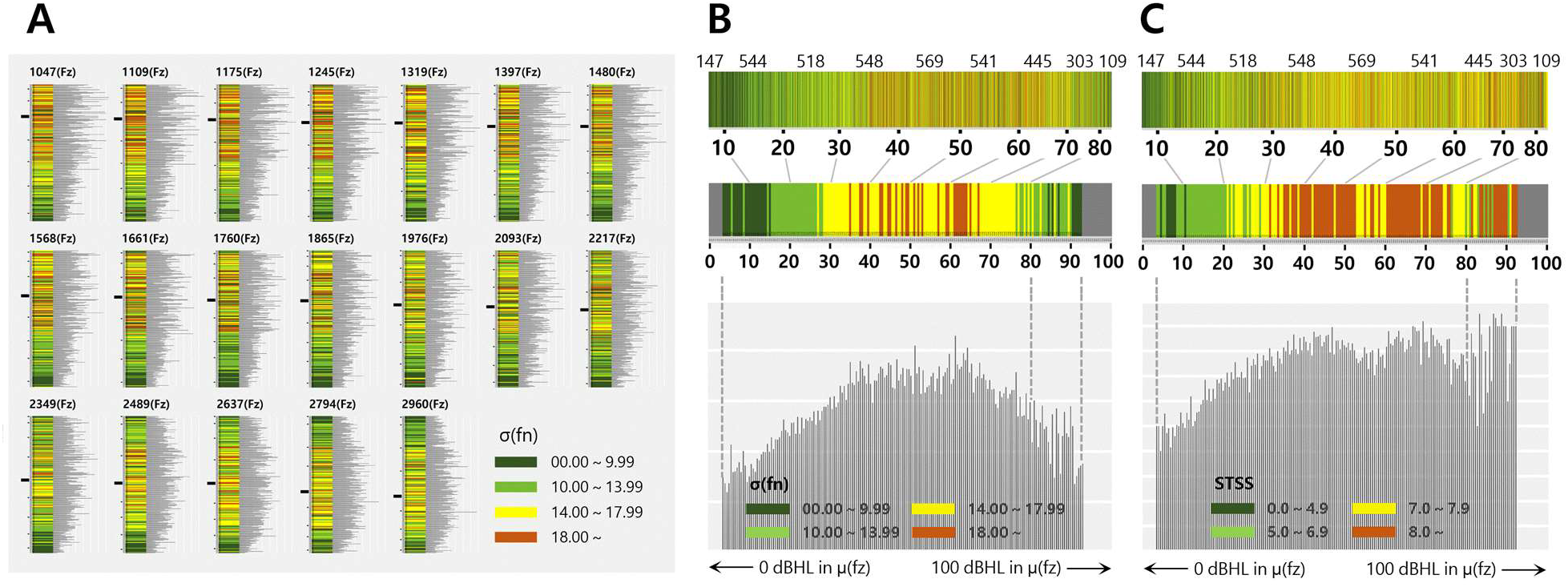
(A) Spectrogram-like representation of σ(f_n_) levels (N = 196) in each of 19 harmonic templates. (B) Summed representation of σ(f_n_) (N = 3,724) from 196 individual σ(f_n_) levels at the 19 harmonic templates and the normalized representation (N = 179) with equivalent intervals in 0.5-dB steps. (C) Summed representation of STSS (N = 3,724) from the 196 individual STSS levels at the 19 harmonic templates and the normalized representation (N = 179) with equivalent intervals in 0.5-dB steps.

### Gaussian shapes of σ(f_n_) and tinnitus severity against μ(fz)

The level of σ(f_n_) in a certain Fz was deduced from the distance between the auditory thresholds of individual harmonics sharing a common fundamental frequency. Being peaked around 01(Fz) in the X-axis, the 50~60 dBHL in the Y-axis, and the 15~20 dBzHL in the Z-axis, our results showed that the distribution of the σ(f_n_) levels (N = 3,724) from 196 chronic tinnitus patients tended to follow a Gaussian curvature across the X and Y-axes, which represented the Fz fundamental frequency (f_0_) and the μ(fz) of Fz, respectively. We assumed that the σ(f_n_) levels on the Z-axis for dBzHLs from chronic tinnitus patients occupied the middle area in the Gaussian curve against the μ(fz)-axis for dBHLs and a right-half Gaussian curve against the fundamental frequency of the 19 harmonic templates in Hz. Figure 5A shows that the average distribution of σ(f_n_) from the 196 chronic tinnitus patients followed the middle area of a Gaussian curve against μ(fz) (*p* = 0.75912, Lilliefors test of normality). Figure 5B shows that the average distribution of σ(f_n_) from the 19 harmonic templates followed a right-half Gaussian curve against the fundamental frequency of the 19 harmonic templates (*p* = 0.24901, Lilliefors test of normality). The 3,724 samples of σ(f_n_) with f_n_ (n = 0, 1, 2, 3) truncated at the fourth harmonics of Fz were normalized with 179 grids in 0.5-dB steps on the Y-axis. As shown in Figure 5C, the relationship between 179 μ(fz) values and 179 STSS levels in the normalized representation in 0.5-dB steps was positive, with a correlation coefficient of 0.69 (*p* = 0.00000). If the analysis was limited to μ(fz) values below the 80 dBHL, the correlation coefficient was 0.91 (*p* = 0.00000). As for fluctuation patterns above the 80 dBHL, a positive relationship was still observed between the fluctuation patterns in both σ(f_n_) and STSS (*r* = 0.4, *p* = 0.04816), as shown in Figure 5D.

**Figure 5.**
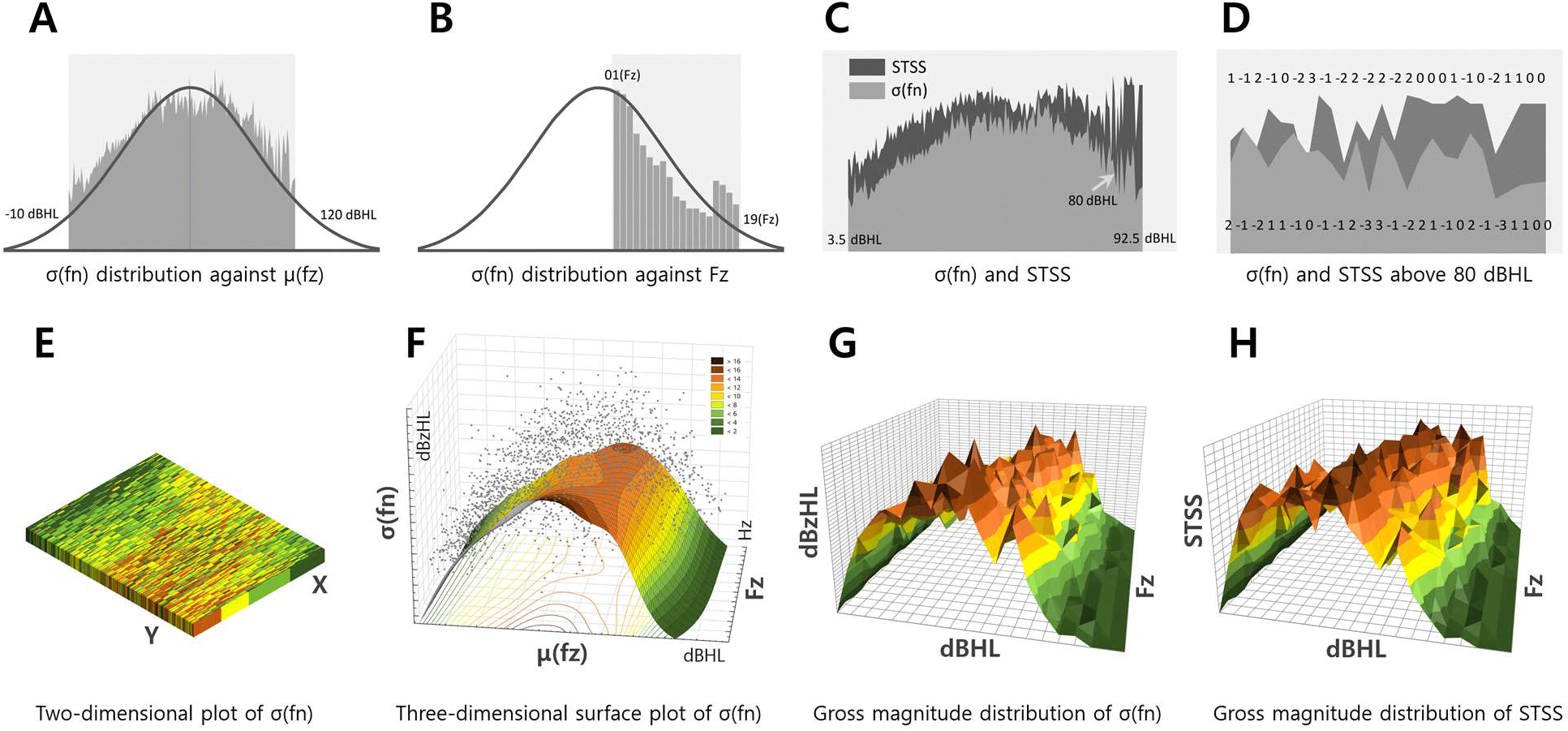
(A) Gaussian curve and distribution of average levels of σ(f_n_) against μ(fz). Lilliefors test confirmed that the distribution of σ(f_n_) fit with the middle part of the Gaussian curve (*p* = 0.75912). (B) Gaussian curve and distribution of the average levels of σ(f_n_) against the 19 fundamental frequencies of Fz. Lilliefors test confirmed that the distribution of σ(f_n_) fit with a right-half Gaussian curve (*p* = 0.24901). (C) Normalized representation (N = 179) based on equivalent intervals in 0.5-dB steps from the 3.5 dBHL to the 92.5 dBHL on the μ(fz)-axis. The two parameters of σ(f_n_) and STSS showed a strong positive correlation (*r* = 0.91) from the 3.5 dBHL to the 79.5 dBHL on the μ(fz)-axis. (D) A moderate positive correlation (*r* = 0.4) was found in fluctuation patterns between the two parameters of σ(f_n_) and STSS from the 80 dBHL to the 92.5 dBHL on the μ(fz)-axis. (E) A 2D representation of individual σ(f_n_) levels (N = 3,724) against μ(fz) and the 19 harmonic templates in 196 chronic tinnitus patients. This was a unified matrix of Figure 4A and Table 2 without normalization by equivalent dB intervals on the Y-axis. (F) A 3D surface plot against Figure 5E by distance-weighted least-squares with normalization in 0.5-dB steps on the Y-axis. (G) Gross magnitude distribution of σ(f_n_) levels (N = 3,724) in dBzHL against equivalent 5-dB intervals on the Y-axis in dBHL and the 19 fundamental frequencies of Fz on the X-axis in Hz. (H) Gross magnitude distribution of STSS by 5-dB steps.

Figure 5F shows a surface plot of σ(f_n_) levels on the Z-axis in dBzHL which represents the average distribution of 3,724 σ(f_n_) levels against the μ(fz)-axis in dBHL with the normalization of equal intervals in 0.5-dB steps and against the X-axis of the 19 fundamental frequencies of Fz in Hz. This was a 3D representation by distance-weighted least-squares against Figure 5E, which is a unified 2D representation of the 3,724 σ(f_n_) levels of the 19 harmonic templates shown in Figure 4A and Table 2 without normalization by equivalent dB intervals one the Y-axis. The gross magnitude distribution of 3,724 σ(f_n_) levels in the 196 chronic tinnitus patients is shown in Figure 5G with equivalent 5-dB intervals on the Y-axis in dBHL and the 19 fundamental frequencies of Fz on the X-axis in Hz. The gross magnitude of the auditory instability in each harmonic template was calculated by the formula μ(σ(f_n_)) * N/19.6 with 3,724 σ(f_n_) levels from 196 chronic tinnitus patients. The formula was devised to quantify the gross magnitude (in dBzHL) of auditory instability σ(f_n_) in 196 chronic tinnitus patients against Fz in Hz and μ(fz) in dBHL. Figure 5H shows the gross magnitude distribution in STSS evaluated using the formula μ(STSS) * N/19.6. There was a strong positive correlation between the gross magnitude distributions of σ(f_n_) and STSS (*r* = 0.97, *p* = 0.00000).

## Discussion

Neural studies on tinnitus have shown that tinnitus is due to hearing impairment, i.e., decreased auditory input. However, there are clinical reports on severe tinnitus patients with normal audiograms. First, we reconsidered the most common consensus on the relationship between decreased auditory input and the severity of tinnitus. There are still no systematic data on the exact relationship between hearing impairment and tinnitus loudness, although recent findings^20, 21^ have pointed out a nonlinear relationship between hearing impairment and tinnitus. A few research studies have provided basic perspectives on the correlation between hearing loss and the severity of tinnitus^1, 22, 23, 24^. Their conclusions, however, do not guarantee that the relationship between decreased auditory input and tinnitus severity is linear. While one trend is moving toward studies on the neural mechanisms of tinnitus, the other argues behavioral aspects of severe tinnitus in normal audiograms. This argument starts with existing octave-based traditional audiometry in which the audiometric resolution is too low to see the fine structure in behavioral hearing thresholds. The answer to the question of why there exists severe tinnitus with normal hearing is in the answer to another question about what “normal” means. There might be numerous cases of notched hearing loss at frequencies not being tested by traditional audiometry. The auditory instability of harmonic templates has never been accounted for in current tinnitus studies. If we can accept a paradigm shift from traditional PTA to algorithmic PTA, one-bark resolution audiometry at least will be required for normal hearing or mild hearing loss patients who experience chronic tinnitus. As shown in Figure 6, one of our raw data showed why the current audiometric definition of “normal” should be revisited.

**Figure 6.**
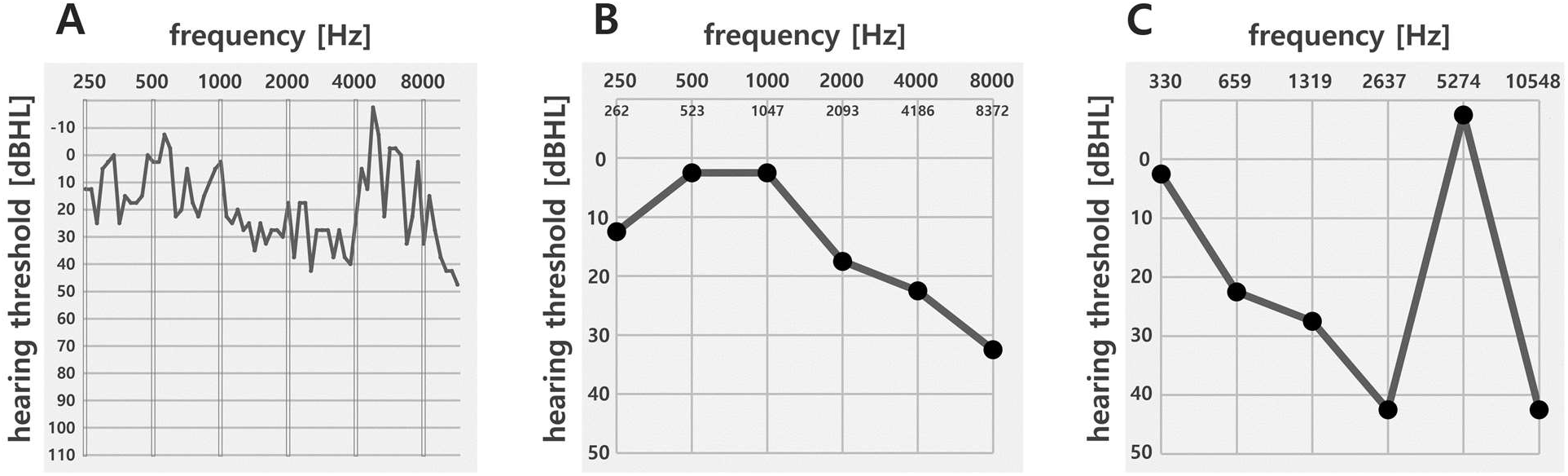
Comparison between conventional audiogram and 1~1.5 bark-shifted audiogram obtained from a chronic tinnitus patient (#1810012) who showed auditory instability levels up to 16 dBzHL with an STSS of 8. While #1810012’s hearing was normal according to the conventional averaging method in PTA, the 1~1.5 bark-shifted audiogram revealed that #1810012’s hearing was not normal. (A) Threshold fine structure in 67 frequency bands with 1/12 octave resolution in patient #1810012. (B) Conventional audiogram of #1810012. (C) 1~1.5 bark-shifted audiogram of #1810012.

Our computational model predicted that the correlation between decreased auditory input and tinnitus severity followed a left-half Gaussian curve up to a plateau around the 15~20 dBzHL in σ(f_n_) against about a 50~60 dBHL in μ(fz). It seemed that the decreased auditory input could be the origin of the increased severity of tinnitus at least up to either the peak or the plateau of a Gaussian curve where the most unstable neural phasing happened in the microsecond range affiliated with the degraded temporal capacity probably due to a malfunction in synaptic gating. Even if the hearing threshold is in the normal range assessed by traditional PTA, there could be a severe disparity between different harmonic components in a certain harmonic template. Neither auditory instability as a function of σ(f_n_) in the dBzHL nor auditory capacity as a function of μ(fz) in the dBHL could have been in the scope of clinical research to date since most behavioral studies on auditory thresholds were performed in clinical practice using octave-based PTA. As shown in Figure 7B, 18 audiograms from normal-hearing or mild hearing loss chronic tinnitus patients revealed transient threshold fluctuations due to burst firing and auditory neural synchrony at deprived frequency regions^25, 26^. A high-resolution audiogram by algorithmic PTA provides precision sources to evaluate both the auditory capacity in dBHL and the auditory instability in dBzHL in a certain harmonic template. It seems that octave-based audiometry is not a proper tool to measure the behavioral hearing thresholds of normal-hearing or mild hearing loss since most patients with normal-hearing or mild hearing loss still have relatively excellent frequency specificity in signal reception and its decoding at levels of the cochlea and the central auditory system. The Monte Carlo algorithm as shown in figure 7A enabled chronic tinnitus patients to complete 1/12 octave resolution audiometry from 250 Hz to 12,000 Hz in both ears within 15~20 minutes. The average turnaround time for the 43 frequencies from 1,047 Hz to 11,840 Hz was 10 minutes in the 196 tinnitus patients in the present study. The Monte Carlo method in audiometry is analogous to randomized Bekesy audiometry. To project the tinnitus spectrum in a 3D Cartesian coordinate, 67 test signals in 5-dB steps are presented as random functions of frequency and intensity. Figure 7B is a 2D representation of various hearing threshold fine structures with 1/12 octave frequency resolution on the conventional audiogram form. Eighteen examples were obtained from normal hearing patients or mild hearing loss patients who experienced chronic tinnitus.

**Figure 7.**
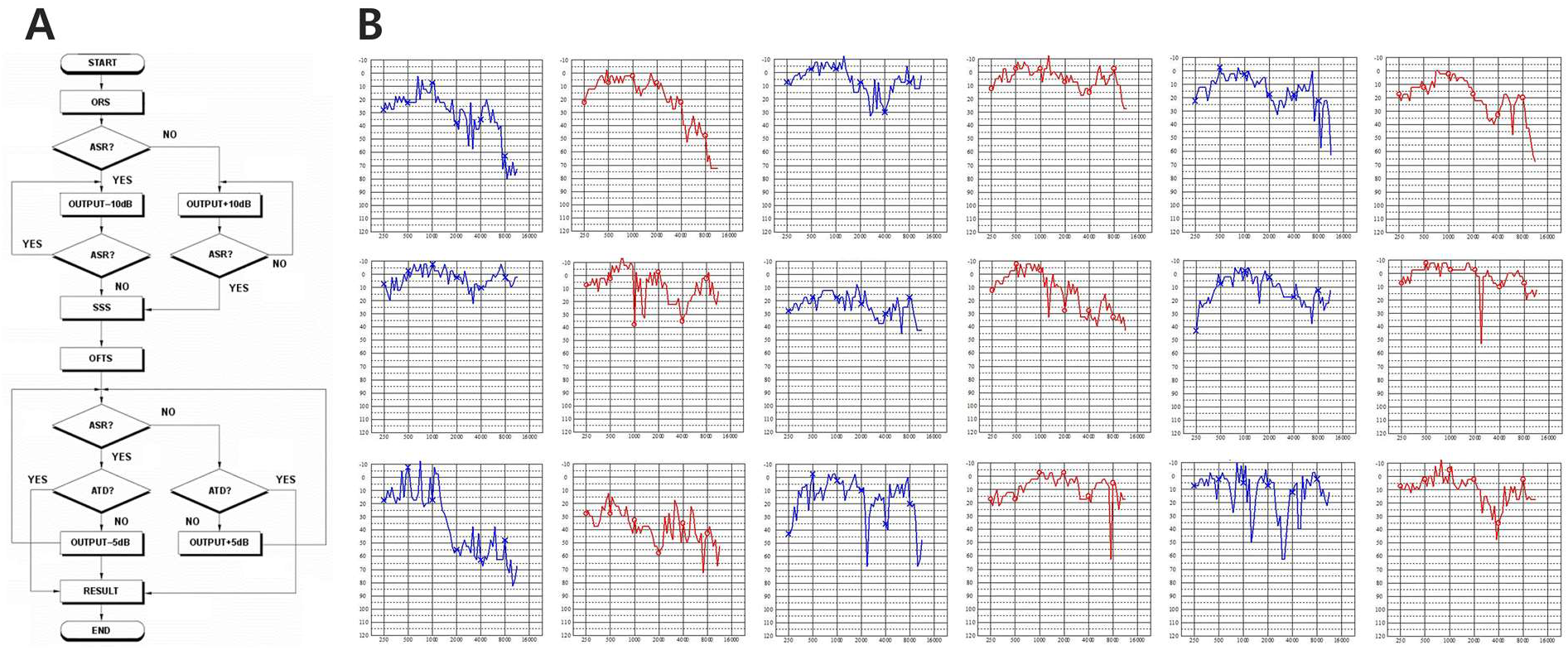
(A) A simple schema of algorithmic pure tone audiometry (PTA) using the Monte Carlo method. (B) Several 1/12 octave frequency resolution audiograms obtained by algorithmic PTA.

The level of auditory instability seems to be closely related to the severity of chronic tinnitus. If it is assumed that the unknown neural impedance (Z) in phasor form is the summation or subtraction of the DC-mediated resistance (R) and the AC-mediated reactance (X), the perceived loudness level of chronic tinnitus may follow a bimodal distribution due to the anomaly of AC/DC-mediated impedance fluctuation. From the peak or the plateau of a Gaussian curve where the maximum auditory instability arises from the most unstable neural phasing to the right tail where AC/DC-mediated underdamped oscillation or fluctuation might intervene, especially from severe hearing loss regions above 80 dBHL in μ(fz), the level of auditory instability will move with the right-half trajectory of a Gaussian trajectory toward the most stabilized passive zero at which the severity level of tinnitus fades completely at the end of the profound hearing loss regions i.e., the DC-dominant passive auditory unit. Although clarifying the unknown neural AC/DC impedance in active harmonic templates was beyond the scope of the present study, it seems that the magnitude of the auditory neural impedance Z in phasor form might be a summation/subtraction of the biological resistance R in a certain auditory unit and neurological reactance X in the active auditory system defined as Z = R ± jX.

We employed a lowercase letter z to indicate complex impedance in the active harmonic templates. As shown in Figure 8, a Rayleigh curve with horizontal flipping, called a z-weighted Gaussian curve in the present study, represented the trajectory of loudness level in chronic tinnitus against the μ(fz)-axis for auditory capacity, especially in the right-half region. The Gaussian model was fit to both the distributions of auditory instability σ(f_n_) and the loudness level of simple tinnitus against the μ(fz)-axis in the active harmonic templates. A sinusoidal curve is assumed to estimate the physical amplitude level of simple tinnitus in a certain passive auditory unit.

**Figure 8.**
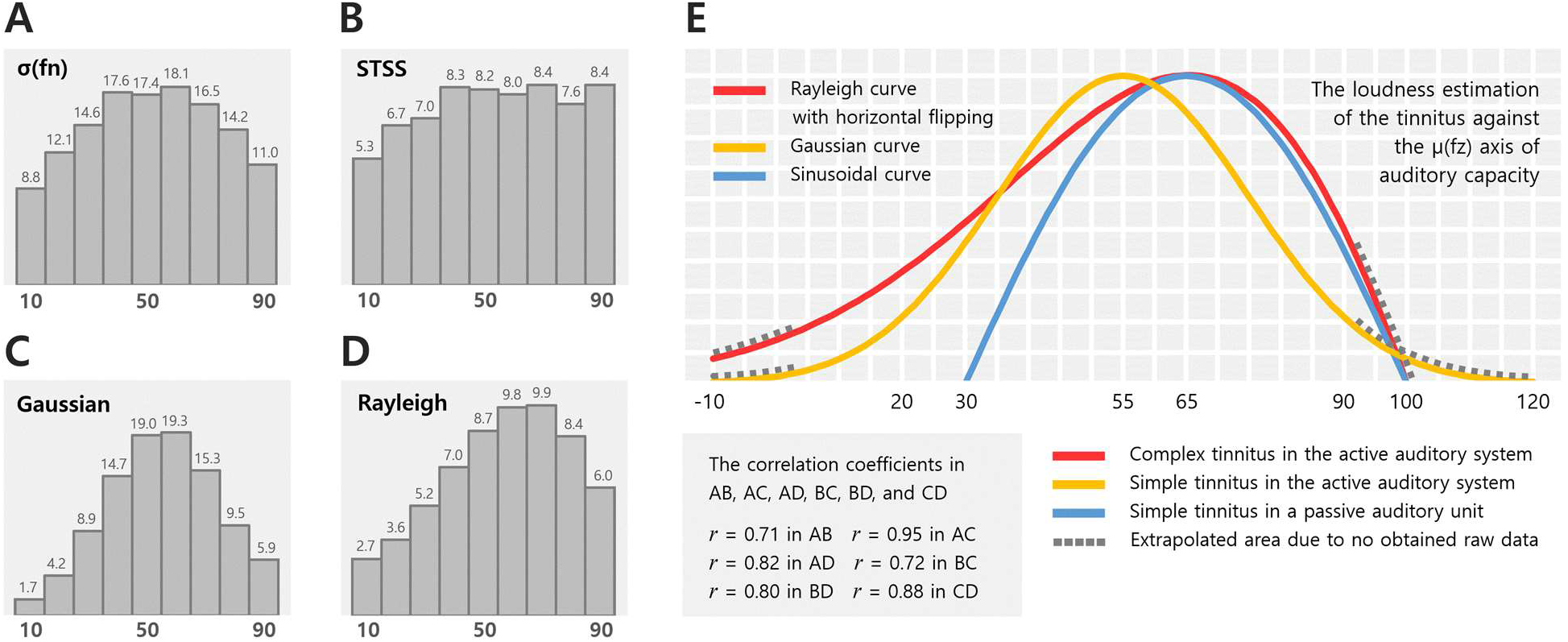
Histograms of the average distribution of σ(f_n_) and STSS against μ(fz) in 196 chronic tinnitus patients and those simulated by the models. (A) Histogram of the average distribution of auditory instability σ(f_n_) (N = 3,724) in chronic tinnitus patients (N = 196) in 10-dB steps. (B) Histogram of the average distribution of simplified tinnitus severity score (N = 3,724) in chronic tinnitus patients (N = 196) in 10-dB steps. (C) Histogram simulated by the Gaussian model without z-weighting. (D) Histogram simulated by the z-weighted Gaussian model, i.e., the Rayleigh model with horizontal flipping. (E) Three models to estimate tinnitus loudness against auditory capacity.

When chronic tinnitus patients participate in the tinnitus matching test and surveys of tinnitus severity, they are required to have their own capabilities to assess not only the pitch but also timbre information. The complex spectra in chronic tinnitus make it difficult to conduct exact assessments of pitch, loudness, and timbre in the tinnitus matching tasks. For instance, if a patient has complex tonal tinnitus containing octave (1:2 ratio)-related tinnitus components, e.g., both 4 kHz and 8 kHz, he or she cannot easily perceive the partial pitches of tinnitus because spectral fusion occurs in the same harmonic template^27^. Most chronic tinnitus patients have higher complexity in their tinnitus spectra than mild or simple tinnitus patients. Mild or simple tinnitus patients tend to quickly identify the pitch and timbre of their tinnitus.

Algorithmic PTA enables a stochastic approach to tinnitus severity evaluation. We assumed that the loudness of chronic tinnitus followed a vector-valued function using a Gaussian integral. As shown in Figure 4B, the distribution of σ(f_n_) levels in the 196 chronic patients had a Gaussian trajectory. The STSS distribution shown in Figure 4C, however, seems a bit right-skewed. The Rayleigh curve with horizontal flipping was the same as the curve induced by the z-weighted Gaussian integral in this study. The distribution of tinnitus loudness was evaluated by vector integral calculus using a formula (3). We considered vector 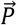 as the length of the line from P(0, 0) to P(y, z), which could be any point on the Gaussian trajectory defined as

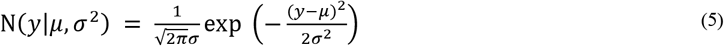

where *y* is the auditory capacity μ(fz). The Gaussian integral of vector 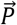

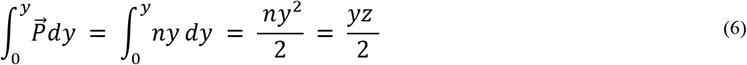

corresponds to the triangle area where the function 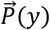 is *ny*, *y* is the base of the triangle representing the auditory capacity μ(fz), and *z* is the height of the triangle representing auditory instability σ(f_n_). If we consider coefficient *k* as the peak or plateau of the Gaussian trajectory, then the active trajectory *L_c_* of the loudness of chronic tinnitus in the active harmonic templates is defined as either

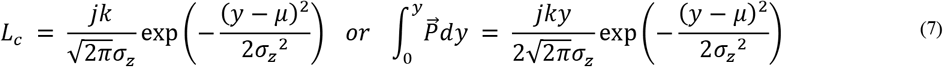

where σ_z_ is the z-weighted standard deviation of the normal Gaussian distribution, weight z is 1.3 which is the coefficient evaluated in this study, and *j* is a proportional factor considered to be 2.159. If the evaluated value of *k* is approximately 20 according to Figure 4B, then *Lc* is

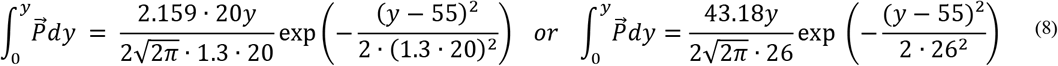

where the standard deviation of the Gaussian distribution is 20 and the mean *μ* of y is 55. The type of formula (8) is the same as that of formula (3).

The horizontally flipped curve of a normal Rayleigh distribution shown in Figure 8E is defined as

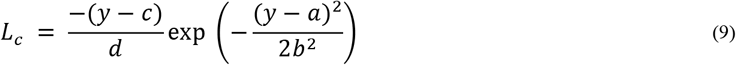

where *a* is 120, *b* is 43.5, *c* is 100 and *d* is *π*/4.

To project a 3D representation of the loudness of chronic complex tinnitus, formula (4) is evaluated with the data of Figure 4C, Figure 5H, and Figure 8E given by either

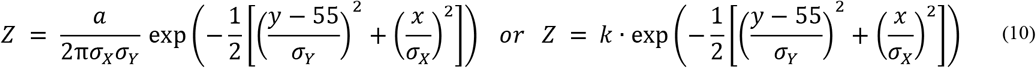

where the capital letter *Z* was employed to indicate the perceived loudness level of chronic complex tinnitus. We considered the coefficient *k* as the peak or plateau in both Gaussian trajectories against the Fz-axis in Hz and the μ(fz)-axis in dBHL. If *σ_x_*, which indicates the standard deviation of a Gaussian curve against Fz is 10, if *σ_Y_*, indicating the standard deviation of a Gaussian curvature against μ(fz) is 20, and if coefficient *k* is 20, then formula (10) can be expressed as:

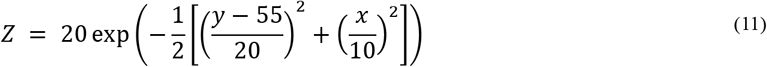

where *x* ≥ 0.

A better temporal resolution in a certain harmonic template (Fz) allows higher resolvability for multiple harmonic inputs^28, 29^. An unstable temporal fine structure in a certain Fz may facilitate the degeneration of auditory capacity by dephasing or defocusing at the neuronal synaptic level in the microsecond time scale^30, 31^. Up to the peak or plateau on the z-weighted Gaussian curve, the instability of harmonic templates resulted in an increased failure rate in auditory phase locking which might be a key origin of the higher severity of chronic tinnitus up to around the 65 dBHL threshold. The underlying mechanisms of the degraded success rate of phase locking may be explained by several recent studies on the randomness in the sequence and timing of reactions^32^, the degree of molecule fluctuations due to stochastic defocusing^33^, and the transient phase gradients evoked by stochastic dephasing^34^. An intriguing aspect in behavioral STSS levels above the 80 dBHL in the μ(fz)-axis was that fluctuation patterns in the STSS levels from the 80 to the 92.5 dBHL showed synchronized shapes with instability patterns of σ(f_n_). Furthermore, the amplified dynamic range in the STSS levels occupied a more enlarged area than that of σ(f_n_) with increases in the degeneracy in μ(fz). With regard to this enlarged convexity in the STSS levels, we assume that the amplified and fluctuated pattern in the severity of chronic tinnitus above the 80 dBHL in μ(fz) will follow an exponential bistable transfer function at the saturated mean-spike-rate level in the aged harmonic template, which is the aged phase locking processor, as a weak signal processor or an underdamped bistable system^2,3, 35, 36, 37, 38,39^. The enlarged convexity in STSS levels tends to initiate at the 50~60 dBHL in μ(fz), as shown in Figures 4C and 5C. Further studies on vector strength in neural phasing are needed to investigate any powers related to synaptic gating control and lowpass filtering above the 50~60 dBHL in μ(fz)^3, 40^.

Along with the precise quantification of chronic tinnitus, one of the aims of the present study was to find a possible way to stabilize and improve any abnormalities in neural phasing in the active auditory system. As for the reversibility of auditory capacity, there are a few recent studies on the correlation between behavioral modality and outcomes in an active neural network. One study reported that an increased listening effort could change functional connectivity in the central nervous system^41^. Another study focused on auditory threshold amelioration by an attention-free listening effort^42^. Although the behavioral modality in both studies was an active listening effort, the latter is not attention-involved. The method of threshold sound conditioning (TSC) uses bistable auditory streaming conducted at a barely audible level with attention-free awareness. It seems that threshold stimulation can influence dynamic bistability during the neural phasing process. The phase in the dynamic bistable circuit (i.e., the temporal fine structure of human active harmonic templates) might be reformed by auditory threshold conditioning, thereby changing the auditory capacity. Evidences have shown that changes in the overall firing of neural ensembles are led by stimulus-induced phase resetting^43, 44^. TSC differs from other existing methods using bistable auditory streaming conducted at audible levels^45^. It is noticeable that two different studies on hearing threshold improvement using a notched sound^46^ and a threshold sound^42^ reported similar findings. The experimental results in both studies might have the same substrate on which stochastic impedance levels in deprived frequency regions may become a robust biomarker for both the reversibility of sensorineural hearing loss and the reducibility of tinnitus loudness.

A recent study^5^ reported several cases of tinnitus remission in chronic tinnitus patients. However, that study could not find any clear factors to explain why tinnitus resolved in those patients. The proposed z-weighted Gaussian model in the present study could easily explain the reason why those patients could achieve tinnitus remission. We found that behavioral hearing thresholds around their salient tinnitus frequencies were in the region of either normal hearing or severe hearing loss. Both hearing levels were located in each tail of a Gaussian curve representing the σ(f_n_) level distribution against μ(fz) where their unstable harmonic templates could easily achieve a completely stable condition. Various therapies have been adopted in tinnitus treatment in clinical practice over the last decades. However, attention has not been paid to positive or negative changes in patients’ hearing threshold fine structures during or after the treatment of chronic tinnitus in clinics. Our computational models and analysis of the behavioral hearing threshold fine structure demonstrated that tinnitus remission during or after treatment may involve more degraded fine hearing structures. In other words, tinnitus relief by listening to any audible acoustic signals should be secured from the unwanted acceleration of degeneration in the auditory temporal fine structure.

The frequency specificity in the human auditory system seems to have its physiologic roots in the very finegrained tonotopic map in the cochlea^12, 13, 47^ which is not only an entrance for external vibration toward the central auditory system but also an exit through the efferent auditory pathway for dephasing artifacts induced by phase instability from stochastic fluctuation and perturbation in the microsecond time domain^34^. It is plausible that tinnitus remission can be achieved by improved frequency specificity in each partial of the harmonic templates without any significant deterioration in auditory capacity. We conclude that auditory instability in the harmonic templates is a key factor in increasing the loudness level of chronic tinnitus, especially in both complex and combined types of tinnitus. The higher loudness level in chronic tinnitus seems to be from the lower success rate of auditory phase locking due to unstable harmonic templates. Further studies are needed to investigate a useful method for stabilizing the auditory neural network without any deterioration in the hearing threshold fine structure in the harmonic templates.

## Data Availability

The data and code that support the findings of this study are available from the corresponding author on reasonable request.

## Acknowledgments

The authors thank 17 clinics for helping with algorithmic PTA data acquisition.

## Author Contributions

S.K and D.L conceptualized and designed the study and drafted the initial manuscript. S.K drafted the initial manuscript. D.L carried out the mathematical modeling. S.J reviewed the manuscript and gave critical feedback during the elaboration of the manuscript. Songwha K conducted data acquisition and statistical analysis. Sunghwan K reviewed the manuscript and gave critical feedback during the elaboration of the manuscript. W.D developed the algorithms for computational modeling of this study. E.K reviewed, revised, and approved the final manuscript as submitted.

## Notes

### Competing Interest Statement

Sangyeop Kwak is the Chief Science Officer of Sound Vaccine, Inc. and AudioCardio, Inc. He also holds intellectual property ownership over the algorithmic PTA described in this study. Daehee Lee is the Chief Technology Officer at Sound Vaccine, Inc. Sungshin Jang is the Head Researcher at Sound Vaccine, Inc. Songhwa Kim is the Senior Researcher at Sound Vaccine, Inc. Sunghwan Kim is the Assistant Researcher at Sound Vaccine, Inc. Woojin Doo is the Principal Researcher at Sound Vaccine, Inc. Eunyee Kwak is the Chief Executive Officer for Sound Vaccine, Inc.

